# Elephantid genomes reveal the molecular bases of Woolly Mammoth adaptations to the arctic

**DOI:** 10.1101/018366

**Authors:** Vincent J. Lynch, Oscar C. Bedoya-Reina, Aakrosh Ratan, Michael Sulak, Daniela I. Drautz-Moses, George H. Perry, Webb Miller, Stephan C. Schuster

## Abstract

Woolly mammoths and the living elephants are characterized by major phenotypic differences that allowed them to live in very different environments. To identify the genetic changes that underlie the suite of adaptations in woolly mammoths to life in extreme cold, we sequenced the nuclear genome from three Asian elephants and two woolly mammoths, identified and functionally annotated genetic changes unique to the woolly mammoth lineage. We find that genes with mammoth specific amino acid changes are enriched in functions related to circadian biology, skin and hair development and physiology, lipid metabolism, adipose development and physiology, and temperature sensation. Finally we resurrect and functionally test the mammoth and ancestral elephant TRPV3 gene, which encodes a temperature sensitive transient receptor potential (thermoTRP) channel involved in thermal sensation and hair growth, and show that a single mammoth-specific amino acid substitution in an otherwise highly conserved region of the TRPV3 channel strongly affected its temperature sensitivity. Our results have identified a set of genetic changes that likely played important roles in the adaptation of woolly mammoths to life in the high artic.

## INTRODUCTION

Woolly mammoths (*Mammuthus primigenius*), perhaps the most charismatic of the extinct Pleistocene megafauna, have long fascinated humans and become emblems of the ‘ice age’. Unlike the extant elephantids, which live in warm tropical and subtropical habitats, woolly mammoths lived in the extreme cold of the dry steppe-tundra where average winter temperatures ranged from −30° to −50°C (MacDonald et al., 2012). Woolly mammoths evolved a suite of adaptations for artic life, including morphological traits such as small ears and tails to minimize heat loss, a thick layer of subcutaneous fat, long thick fur, and numerous sebaceous glands for insulation (Repin et al., 2004), a large brown-fat deposit behind the neck that may have functioned as a heat source and fat reservoir during winter (Boeskorov et al., 2007; Fisher et al., 2012). They also likely possessed molecular and physiological adaptations in circadian systems (Bloch et al., 2013; Lu et al., 2010) and adipose biology (Liu et al., 2014; Nelson et al., 2014), similar to other artic-adapted species. Mammoths diverged from Asian elephants (*Elephas sp.*) ∼5 MYA (Rohland et al., 2007) and colonized the steppe-tundra 1-2 MYA (Debruyne et al., 2008), suggesting that their suite of cold-adapted traits evolved relatively recently (**Fig. 1**).

**Figure 1.**
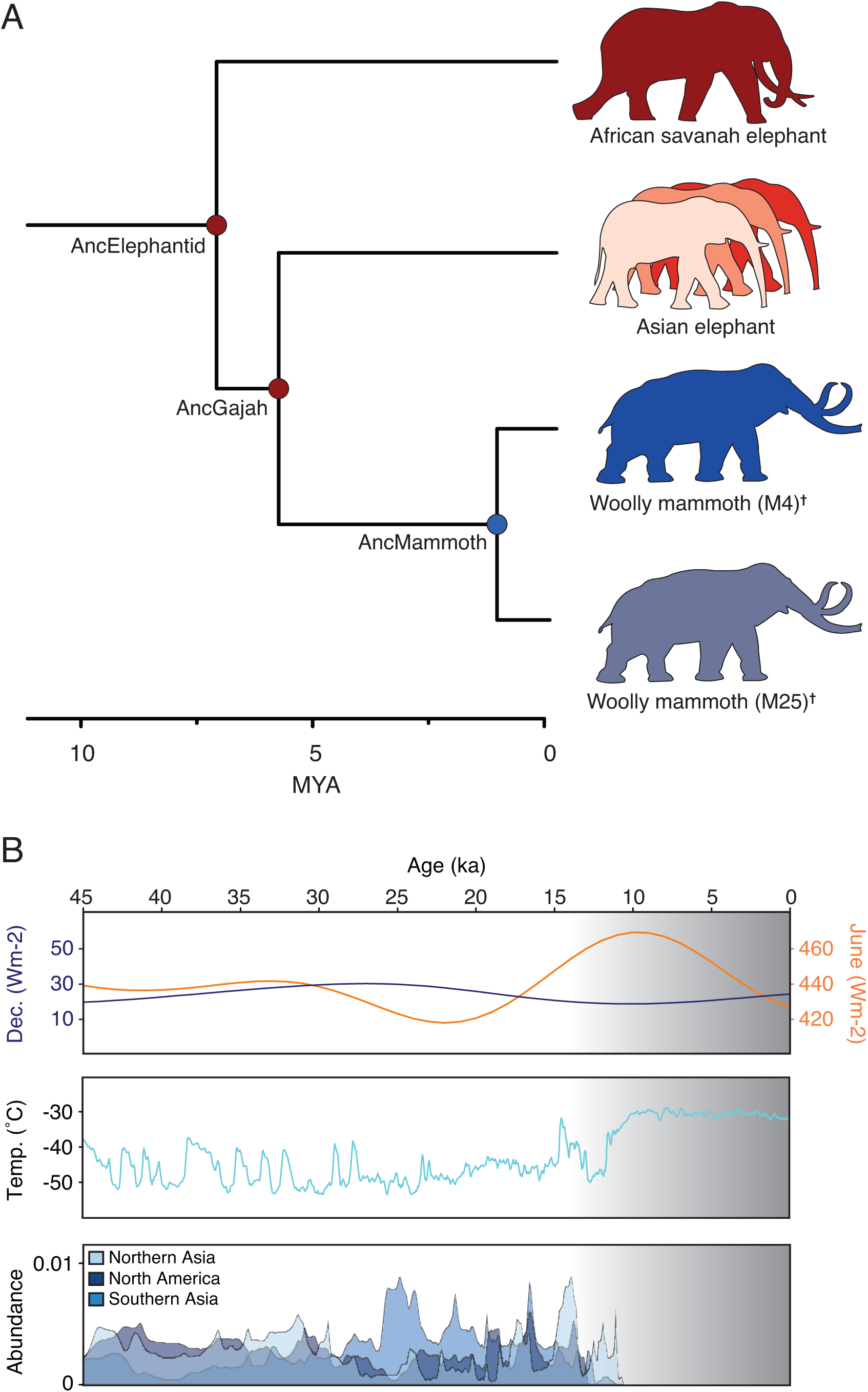
Woolly mammoth phylogeny, ecology, and extinction. (A) Phylogenetic relationships among recent Elephantids. Branches are drawn proportional to time. The ancestor of Asian elephants and mammoths is labeled (AncGajah). (B) Mammoth ecology and extinction. Irradiance at 60°N (upper) in June (orange) and December (blue), arctic surface temperature (middle), and estimated mammoth abundances at three localities (lower) during the last 45 ka years. Data modified from ref. 24.

Identifying the genetic changes that underlie morphological differences between species is daunting, particularly when reconstructing how the genotype-phenotype map diverged in non-model or, especially, extinct organisms. Thus while the molecular bases of some phenotypic traits have been identified, these studies are generally limited to a few well-characterized genes and pathways with relatively simple and direct genotype-phenotype relationships (Chan et al., 2010; Hoekstra et al., 2006; Lang et al., 2012; Smith et al., 2013; Storz et al., 2009). Previous structural and functional studies, for example, have shown that amino acid polymorphisms in the woolly mammoth haemoglobin β/δ fusion gene (*HBB*/*HBD*) reduce oxygen affinity (Campbell et al., 2010; Yuan et al., 2013; 2011) whereas amino acid polymorphisms in both the woolly mammoth and Neandertal melanocortin 1 receptor (*MC1R*) genes were hypomorphic compared to the ancestral alleles (Lalueza-Fox et al., 2007; Römpler et al., 2006). Most traits, however, have complex genotype-phenotype relationships with phenotypic divergence arising through the accumulation of numerous variants of small individual effects rather than one or a few mutations of large effect. Thus candidate gene studies are poorly suited for forward genetic based approaches to trait mapping, and the genetic changes that underlie woolly mammoth adaptations to the artic are almost entirely unknown.

Whole-genome sequencing (WGS) is an invaluable tool for exploring the genetic origins of phenotypic differences between species because one can identify fixed and polymorphic variants across the genome without respect to *a priori* defined genes and pathways. However, distinguishing functional from nonfunctional variants in WGS data can be difficult (Cooper and Shendure, 2011). To determine genetic changes that underlie cold-adapted traits in woolly mammoths we sequenced the genomes of three Asian elephants and two woolly mammoths to high coverage, and functionally annotated fixed, derived amino acid and loss of function substitutions in woolly mammoths. We found that genes with woolly mammoth-specific substitutions were enriched in functions related to circadian biology, skin, hair, and sebaceous gland development and physiology, lipid metabolism, adipose development and physiology, and temperature sensation. We experimentally validated a fixed woolly mammoth-specific hypomorphic substitution in the temperature sensor TRPV3 that we predict influenced both thermal sensation and hair growth in mammoths. These data provide mechanistic insights into the causes of morphological evolution, and define a set of likely casual variants for future study of woolly mammoth-specific traits.

## RESULTS AND DISCUSSION

### Genome sequencing, assembly, and annotation

We generated Illumina sequence data for two woolly mammoths that died ∼20,000 and ∼60,000 years ago (Gilbert et al., 2008; 2007; Miller et al., 2008), including individuals from the two major lineages of woolly mammoths, clade I (individual M4) and clade II (M25), which diverged ∼1.5 MYA (Miller et al., 2008), and three extant Asian elephants (*Elephas maximus*). We aligned sequencing reads to the genome assembly for the African Savannah elephant (*Loxodonta africana*), resulting in non-redundant average sequence coverage of ∼20-fold for each mammoth and ∼30-fold for each Asian elephant (**SI Fig. 1**).

We identified ∼33 million putative single-nucleotide variants (SNVs) among the three elephantid species (see Methods for details), including ∼1.4 million nucleotide variants fixed for the derived allele in the two mammoths, but for the ancestral allele in the African and Asian elephants. Among the variants were 2,020 mammoth-specific amino-acid substitutions in a total of 1,642 protein-coding genes, including 26 protein-coding genes with premature stop codons (putative loss of function substitutions). We also identified a single gene with a strongly supported, fixed, mammoth-specific copy number increase.

### Functional consequences of woolly mammoth-specific amino acid substitutions

We used several complementary approaches to infer the putative functional consequences of mammoth-specific amino acid substitutions, including classifying substitutions based on their BLOSUM80 exchangeabilities (Henikoff and Henikoff, 1992), predicted functional consequences based on PolyPhen-2 (Adzhubei et al., 2010; 2013), and the inter-species conservation of sites at which substitutions occurred, as w ell as identifying KEGG pathways (Kanehisa and Goto, 2000) and mouse knockout (KO) phenotypes (Blake et al., 2014) enriched among protein-coding genes with fixed, derived amino acid substitutions in the wooly mammoth. Finally we manually selected gene-pathway and gene-phenotype associations for further study according to two criteria: 1) the richness of literature supporting the role of each gene in specific pathways and phenotypes, and 2) the exchangeability, PolyPhen-2 score, and strength of sequence conservation at sites with mammoth-specific substitutions.

We found that genes with fixed, derived woolly mammoth substitutions were enriched for 40 KEGG pathways and 859 mouse KO phenotypes, at an FDR≤0.10 (**Fig. 2A**). Significantly enriched KEGG pathways included ‘circadian rhythm - mammal’ (enrichment [E]=6.71, hypergeometric *P*=2.7×10^−3^, FDR *q*=0.02), ‘fat digestion and absorption’ (E=4.01, hypergeometric *P*=7.9×10^−3^, FDR *q*=0.05), ‘complement and coagulation cascades’ (E=4.28, hypergeometric *P*=5.0×10^−4^, FDR *q*=6.7×10^−3^), and ‘metabolic pathways’ (E=8.39, hypergeometric *P*=2.2×10^−7^, FDR *q*=1.6×10^−5^) (**SI Table 1**). Enriched KO phenotypes included ‘decreased core body temperature’ (E=4.15, hypergeometric *P*=8.0×10^−4^, FDR *q*=7.2×10^−3^), ‘abnormal brown adipose tissue morphology’ (E=2.99, hypergeometric *P*=1.4×10^−4^, FDR *q*=4.0×10^−3^), ‘abnormal thermal nociception’ (E=2.25, hypergeometric *P*=5.4×10^−3^, FDR *q*=0.05), ‘abnormal glucose homeostasis’ (E=1.46, hypergeometric *P*=2.6×10^−3^, FDR *q*=3.2×10^−2^), and many body mass/weight-related phenotypes (**Fig. 2B**).

**Figure 2.**
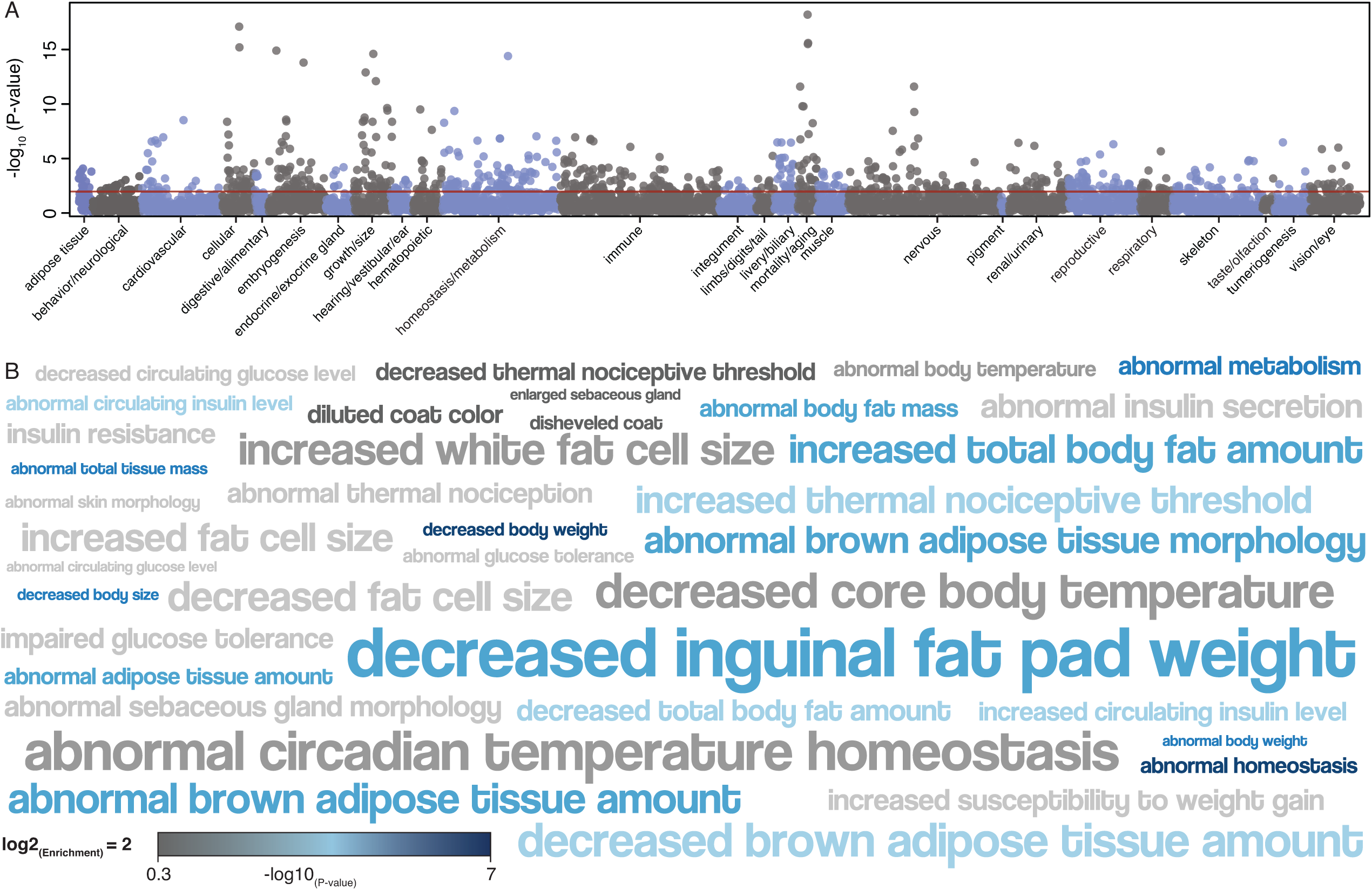
Functional annotation of genes with Woolly Mammoth specific amino acid substitutions. (A) Manhattan plot of mouse knockout phenotypes enriched among genes with fixed, derived mammoth amino acid changes. −log10(hypergeometric P-values) are shown for each phenotype, phenotypes are grouped by anatomical system effected. Horizontal red lines indicates FDR=0.1. 551 genes with fixed, derived mammoth amino acid changes have mouse knockout data. (B) Word cloud of 40 selected mouse knockout phenotypes enriched among the protein coding genes with fixed, derived mammoth amino acid changes. Phenotype terms are scaled to the log2 enrichment of that phenotype and color coded by -log10 P-value of phenotype enrichment (hypergeometric test).

We also inferred the functional significance of fixed, derived loss of function (LOF) substitutions in woolly mammoth genes. We identified a single KEGG term enriched among the genes with LOF substitutions, ‘fat digestion and absorption’ (E=127.64, hypergeometric *P*=1.0×10^−4^, FDR *q*=1.0×10^−4^), and 48 KO terms enriched among these genes at an FDR≤0.10 (**Fig. 3A**). Enriched KO terms were almost exclusively related to cholesterol, sterol, triglyceride, and lipid homeostasis and metabolism (**Fig. 3B**), such as ‘decreased circulating cholesterol level’ (E=33.15, hypergeometric *P*=5.7×10^−5^, FDR *q*=4.3×10^−3^), ‘decreased sterol level’ (E=30.15, hypergeometric *P*=7.6×10^−5^, FDR *q*=4.3×10^−3^), and ‘abnormal circulating lipid level’ (E=30.15, hypergeometric *P*=7.6×10^−5^, FDR *q*=4.3×10^−3^).

**Figure 3.**
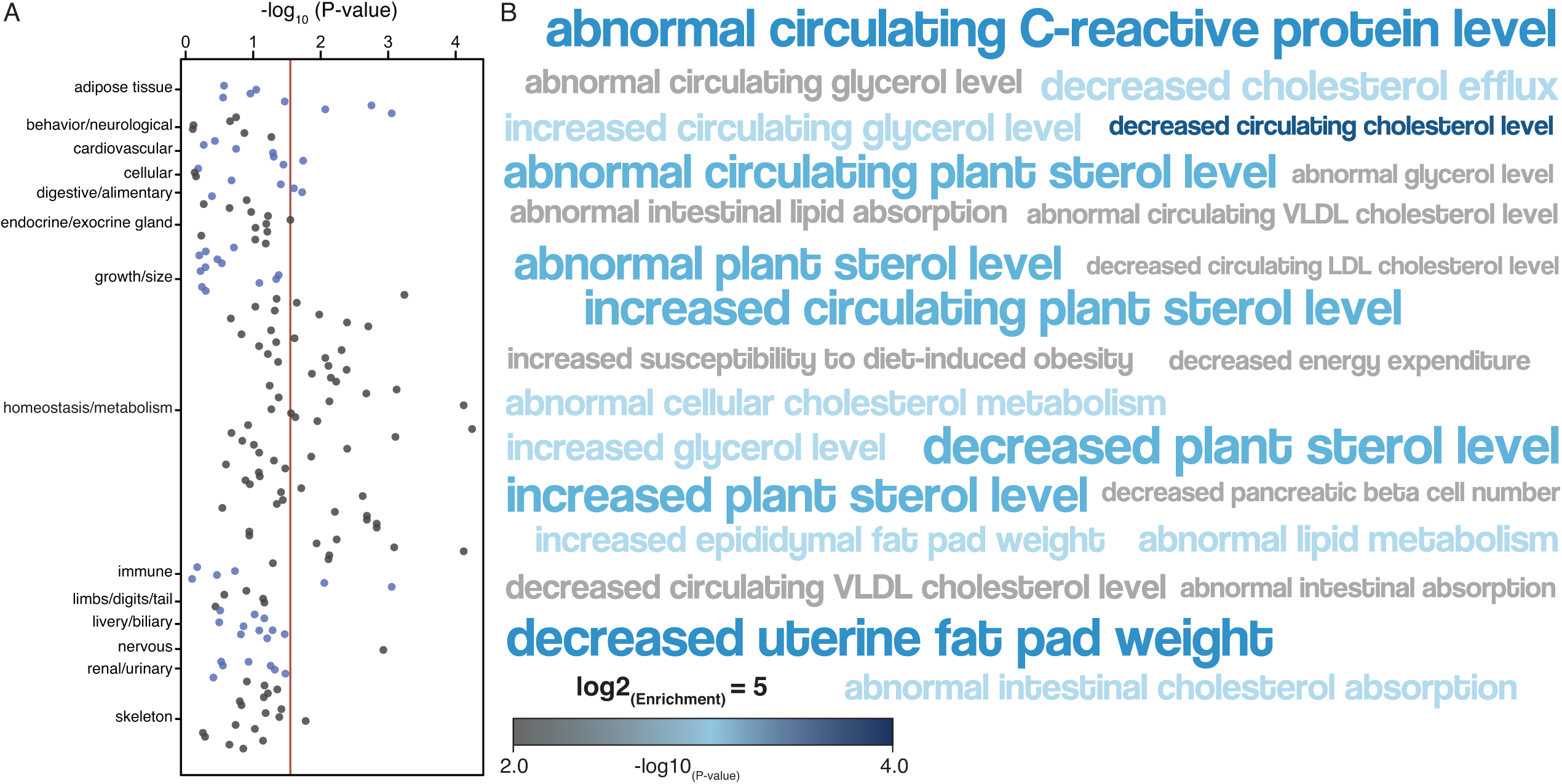
Functional annotation of genes with Woolly Mammoth specific amino loss of function (LOF) substitutions. (A) Manhattan plot of mouse knockout phenotypes enriched among genes with fixed, derived loss premature stop codons in Woolly Mammoths. −log10(hypergeometric P-values) are shown for each phenotype, phenotypes are grouped by anatomical system effected. Horizontal red lines indicates FDR=0.1. (B) Word cloud of 26 selected mouse knockout phenotypes enriched among the protein coding genes with fixed, derived premature stop codons in Woolly Mammoths. Phenotype terms are scaled to the log2 enrichment of that phenotype and color coded by -log10 P-value of phenotype enrichment (hypergeometric test).

### Substitutions in genes associated with circadian biology

Organisms living at high latitudes in the artic experience long periods of darkness during the winter and near constant light in the summer, which prevents polar-adapted species from utilizing daily light-dark cycles to entrain their circadian clocks. Svalbard reindeer (*Rangifer tarandus platyrhynchus*), for example, have lost functioning circadian clocks and circadian rhythmicity in *PER2*, and *BMAL1* (*ARNTL*) expression (Lu et al., 2010). Moreover, several other artic species are also known to have derived circadian systems (Bloch et al., 2013), and we observed that several enriched KO and KEGG terms were related to circadian biology, motivating us to explore circadian genes in greater detail.

Fixed, derived mammoth-specific amino acid substitutions occurred in eight genes associated with circadian biology, including those that play central roles in maintaining normal circadian rhythms and entraining the circadian clock to external stimuli such as temperature. *HRH3*, and *PER2* knockout mice, for example, have abnormal circadian temperature homeostasis (Shiromani et al., 2004; Toyota et al., 2002). *PER2* directly mediates the early adaptive response to shifted temperature cycles and coordinates adaptive thermogenesis by synchronizing *UCP1* expression and activation in brown adipose tissue (Chappuis et al., 2013; Saini et al., 2012). Similarly, neuronal histamine receptors regulate circadian energy homeostasis through *UCP1* expression in brown adipose tissue. *HRH1* KO mice, for example, have abnormal circadian rhythms and abnormal circadian feeding behaviors, including a shift in food consumption from day to night (Inoue et al., 1996). These observations suggest that the circadian system in woolly mammoths may have adapted to the extreme seasonal light-dark oscillations in the high artic.

### Substitutions in genes associated with insulin signaling, lipid metabolism, and adipose biology

The enrichment of genes with derived amino acid (**Fig. 2**) and LOF substitutions (**Fig. 3**) in woolly mammoths that function in lipid metabolism, adipose development, and physiology suggests modifications of these processes may have played an important role in the evolution of woolly mammoths and adaptation to arctic life. We identified 54 genes with fixed, derived amino acid substitutions and KO phenotypes that affect adipose tissue, including phenotypes that alter both the location and abundance of white and brown fat deposits throughout the body. Among the genes with woolly mammoth-specific substitutions were the leptin receptor (*LEPR*), *DLK1* (also known as preadipocyte factor 1) an epidermal growth factor repeat-containing transmembrane protein that regulates adipocyte differentiation, the growth hormone receptor (*GHR*), and Corticotropin-releasing hormone (*CRH*). We also identified 39 genes with KO phenotypes that affect insulin signaling, and found that genes with mammoth-specific amino acid substitutions were enriched in several KO phenotypes related to insulin signaling, including ‘abnormal circulating insulin level’ (E=1.82, hypergeometric *P*=1.0×10^−3^, FDR *q*=1.5×10^−2^), ‘insulin resistance’ (E=2.23, hypergeometric *P*=3.0×10^−3^, FDR *q*=3.5×10^−2^), and ‘impaired glucose tolerance’ (E=1.91, hypergeometric *P*=4.0×10^−3^, FDR *q*=3.7×10^−2^).

### Lineage-specific expansion of the adipocyte regulator *RNASEL* in woolly mammoths

The single well-supported gene duplication we identified in woolly mammoths was *Ribonuclease L* (*RNASEL*), which regulates terminal adipocyte differentiation, lipid storage, and insulin sensitivity (Fabre et al., 2012). We found that the normalized number of reads mapping to the region of the African elephant scaffold 16 encoding the *RNASEL* gene was significantly greater in woolly mammoths than either Asian or African elephants (**Fig. 4A**), consistent with a increase from two to approximately six haploid copies of this gene through relatively recent duplications. We also found that the region of the mammoth *RNASEL* locus homologous to the previously characterized human *RNASEL* enhancer/promoter was included within the duplication, suggesting the duplicate *RNASEL* genes were capable of being actively transcribed. Remarkably, *RNASEL* copy number has expanded specifically in the woolly mammoth lineage but was fixed at the ancestral copy number in the genome of 48 other Eutherian mammals (**Fig. 4B**). *RNASEL*^-/-^ cells, for example, have decreased lipid storage capacity, insulin sensitivity, and glucose uptake, whereas overexpression of *RNASEL* increases lipid storage, insulin sensitivity, and glucose uptake (Fabre et al., 2012). We hypothesize that the increase in copy number of *RNASEL* in woolly mammoths directly affected the development and function of adipocytes.

**Figure 4.**
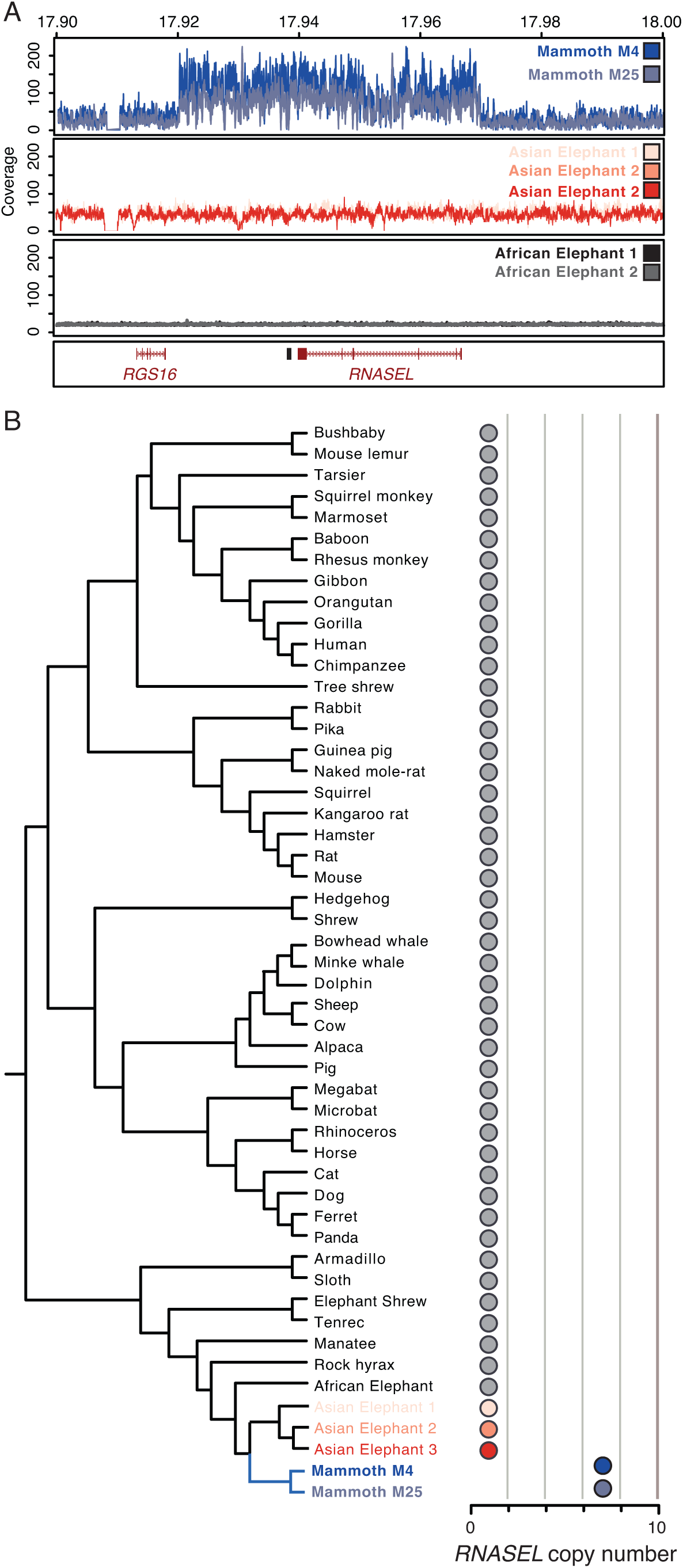
Lineage-specific expansion of the *RNASEL* gene in Woolly Mammoths. (A) Coverage of African elephant (n=2), Asian elephant (n=3), and wooly mammoth (n=2) sequencing reads across the region of African elephant scaffold_16 encoding the *RNASEL* gene. Distances are shown in megabases, the location of the *RNASEL* and *RGS16* genes are shown in red, and the putative *RNASEL* enhancer/promoter in black. (B) *RNASEL* copy number in 49 Eutherian (‘Placental’) mammalian genomes.

### Substitutions in genes associated with skin, hair, and sebaceous gland development and physiology

Like many cold-adapted species, the woolly mammoth coat consisted of two layers – a long, outer layer and a dense inner layer. Unlike other elephants, the skin of mammoths also contained numerous, large sebaceous glands, which are thought to have helped repel water and improve insulation (Repin et al., 2004). Consistent with mammoth-specific amino acid changes contributing to these derived characters, we found that genes with mammoth-specific substitutions were enriched in KO phenotypes related to ‘abnormal sebaceous gland morphology’ (E=2.33, hypergeometric *P*=8.0×10^−3^, FDR *q*=6.3×10^−2^), including substitutions in three genes leading to ‘enlarged sebaceous glands’.

Unlike other elephants, woolly mammoths had thick fur that varied in color from blonde to orange to nearly black (Valente, 1983). Previous studies have found hypomorphic polymorphisms in the woolly mammoth MC1R (Römpler et al., 2006), but these hypomorphic variants may have been relatively rare in some mammoth populations (Workman et al., 2011). Thus variants in other genes may also have contributed to coat color variability in mammoths. We identified 38 genes with mammoth-specific amino acid changes associated with ‘abnormal coat/hair morphology’ in KO mice including derived substitutions in eight genes specifically associated with ‘diluted coat color’. We also found that genes with fixed, derived woolly mammoth substitutions were enriched in ‘hair root sheath’ (hypergeometric *P=*0.006), ‘coat hair follicle’ (hypergeometric *P*=0.013), ‘hair follicle’ (hypergeometric *P=*0.016), skin (hypergeometric *P=*0.018), and ‘hair outer root sheath’ (hypergeometric *P=*0.018). Among the genes associated with dilute coat color is *solute carrier family 7 member 11* (*SLC7A11*), a cystine/glutamate antiporter expressed in melanocytes that regulates pheomelanin (red pigment) synthesis in skin and hair (Chintala et al., 2005). We also identified a woolly mammoth-specific substitution in *LYST*, mutations in which cause dilute hair color and is associated with white fur in polar bears (Liu et al., 2014).

### Substitutions in genes associated with temperature sensation

The most intriguing mouse knockout phenotype enriched among genes with woolly mammoth-specific amino acid changes was ‘abnormal thermal nociception’ (13 genes). For example we identified woolly mammoth-specific amino acid changes in four temperature sensitive transient receptor potential (thermoTRP) channels (**Fig. 5A**), which play central roles in sensing noxious cold (TRPA1 and TRPM4) (Aubdool et al., 2014; Karashima et al., 2009), innocuous warmth (TRPV3 and TRPV4) (Chung et al., 2004; Smith et al., 2002; Xu et al., 2002), and in the vascular response to cold (TRPA1) (Aubdool et al., 2014), as well as PIRT, a small phosphoinositide-binding protein that functions a regulatory subunit of the noxious cold sensors TRPV1 and TRPM8 (Kim et al., 2008; Tang et al., 2013).

**Figure 5.**
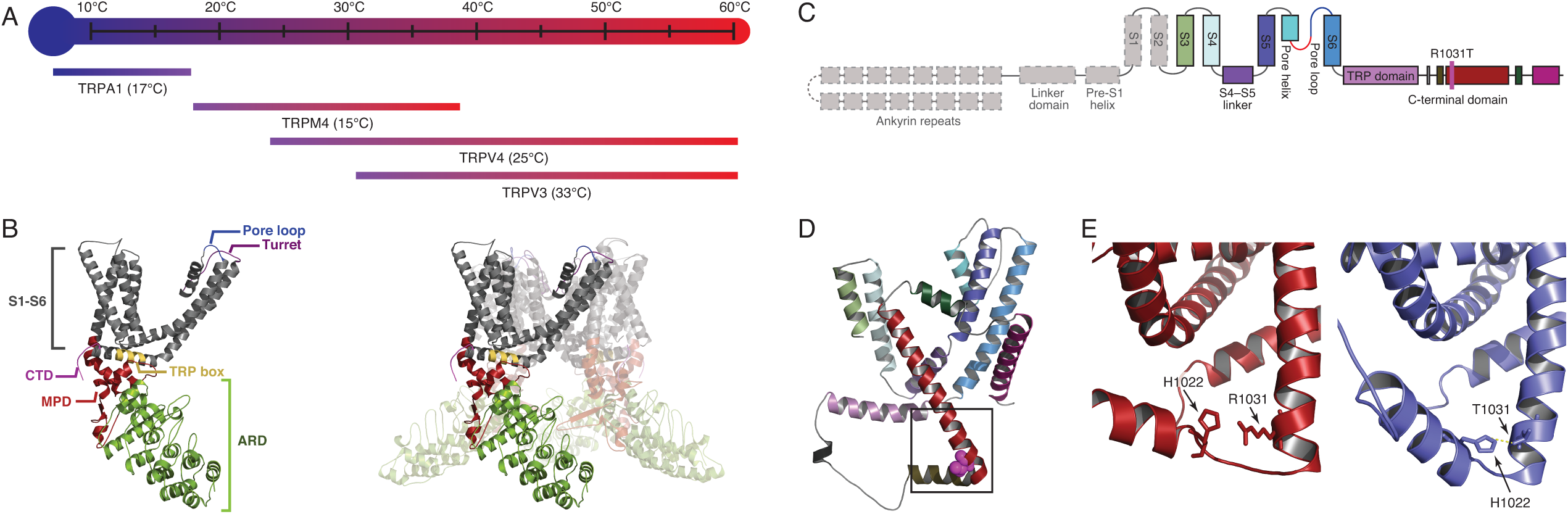
Woolly mammoth-specific amino acid substitutions in thermoTRP temperature sensors. (A) Temperature ranges over which TRPA1, TRPM4, TRPV4, and TRPV3 are active. Threshold temperatures for each channel are shown. (B) Structure of a single TRP subunit (left) and the tetrametic channel (right) viewed from the side. The ankyrin-repeat domain (ARD), transmembrane domains (S1–S6), membrane-proximal domain (MPD), C-terminal domain (CTD), TRP box, pore loop, and pore turret are labeled. Amino acids within the ARD, MPD, pore turret, the outer pore region and in the initial part of S6, the TRP box, and the CTD influence temperature sensing in TRPV and TRPA channels. (C) Diagram of the major structural domains of TRPA1. Grey regions were not included in the TRPA1 structural model. The location of the mammoth-specific R1031T substitution is shown. (D) Cartoon representation of the pore domain of the TRPA1 homology model. The location of the R1031T substitution is shown as magenta colored spheres, helix coloring follows panel (C). (E) Close up of the region boxed in D in the TRPA1 homology model of the AncGajah ancestor (red, left) and Woolly Mammoth (blue, right). The predicted hydrogen bond between T1031 and H1022 in the Woolly Mammoth model is shown as yellow dashed lines.

To infer the putative functional consequences of woolly mammoth-specific amino acid substitutions in thermoTRPs we generated structural models of the ancestral Asian elephant/woolly mammoth (AncGajah; **Fig 1A**) and woolly mammoth (AncMammoth) TRPA1 and TRPV4 proteins, which have well-established roles in thermosensation, based on the recently published high-resolution cryo-EM structure of the TRPV1 channel in the open and closed state (Cao et al., 2013; Liao et al., 2013). We found that the elephantid TRPA1 and TRPV4 proteins were predicted to adopt the common TRPA and TRPV channel structure, which is composed of a series of amino terminal ankyrin repeats (ARD), separated by a membrane proximal domain (MPD) from the six transmembrane helices (S1-S6) that form the ion permeable pore in tetrameric channels (**Fig. 5B**).

We next mapped onto these structure-models the location of the woolly mammoth-specific amino acid changes in TRPA1 (R1031T) and TRPV4 (V658I). Consistent with these substitutions having functional consequences, we found that the TRPA1 R1031T substitution occurred near the end of an alpha helix in the C-terminal domain (**Fig. 5C/D**) and is predicted to introduce a new hydrogen bond between T1031 and the side chain of a neighboring histidine residue (**Fig. 5E**). The mammoth specific TRPV4 V658I substitution occurred at the first site in the S6 helix (**Fig. 6A**), which is part of the outer pore region important for activation of the channel in response to heat (**Fig. 6B**). Indeed, we found that site 658 is located within a cluster of sites that mediate heat activation in the related TRPV3 channel (Grandl et al., 2008) and is homologous to a site in TRPV3 that adopts temperature dependent conformations (**Fig. 6C/D**)(Kim et al., 2013). Site 658 also mediates the interaction between TRPV channels and the agonist vanillotoxin DkTx (**Fig. 6E**) (Cao et al., 2013; Liao et al., 2013). These data suggest the mammoth specific R1031T and V658I substitutions may have affected the gating dynamics in TRPA1 and TRPV4, respectively.

**Figure 6.**
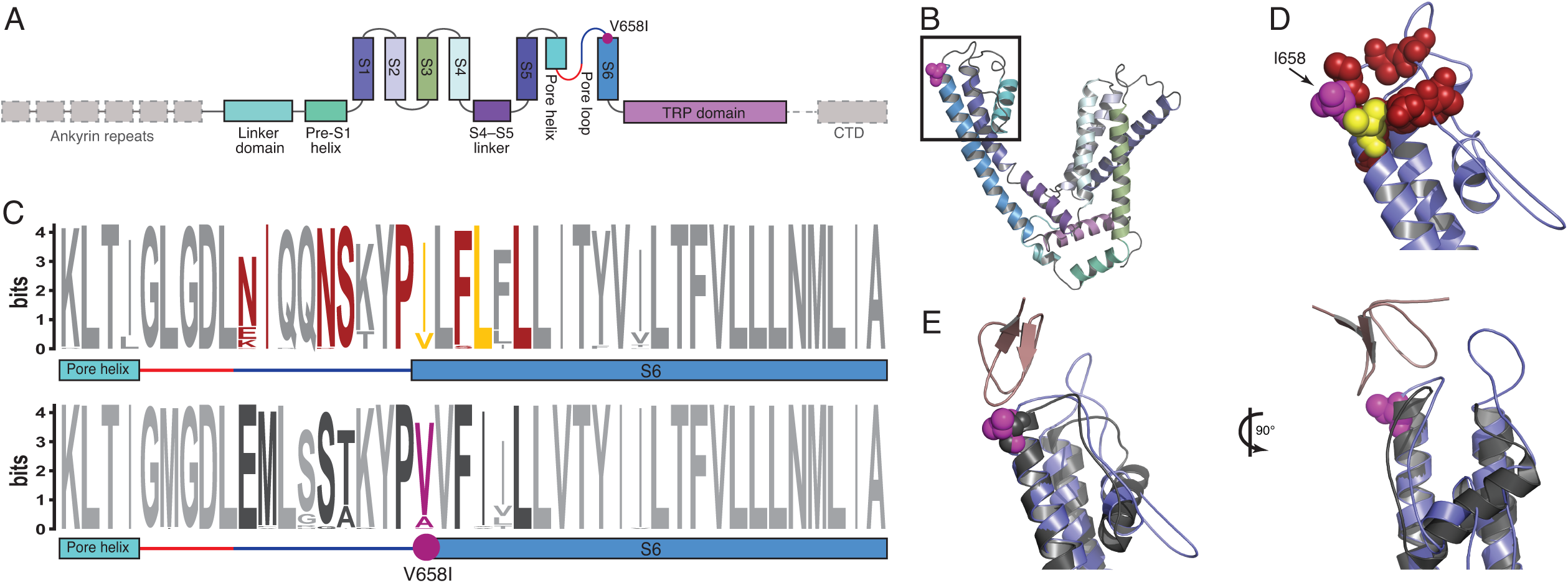
A Woolly mammoth-specific amino acid substitution at a temperature sensitive site in TRPV4. (A) Diagram of the major structural domains of TRPV4. Grey regions were not included in the TRPV4 structural model. The location of the mammoth-specific V658I substitution is shown in magenta. (B) Cartoon representation of the pore domain of the Woolly Mammoth TRPV4 homology model. The location of the V658I substitution is shown as a magenta colored sphere. (C) Conservation of the pore helix-pore loop-S6 region between TRPV3 (upper) and TRPV4 (lower). Residues in TRPV3 that have been experimentally shown to mediate temperature sensing and that have temperature dependent conformations are shown in red and yellow, respectively. Homologous residues in TRPV4 are shown in dark grey, site 658 is shown in magenta. (D) Close up of the region boxed in panel (B). I658 is shown as a magenta colored sphere, homologous residues in mouse TRPV3 that mediate temperature sensitivity are shown as red spheres and residues with temperature dependent conformations in TRPV3 are shown as yellow spheres. (E) Site 658 also mediates the interaction between TRPV channels and the spider vanillotoxin DkTx (pink). I658 is shown as a magenta sphere in the Wooly Mammoth TRPV4 model (blue), the experimentally determined structure of mouse TRPV1 (grey) complexed with DkTx is superimposed onto the mammoth structural model.

### Thermal tuning of the woolly mammoth temperature sensor TRPV3

We found that the mammoth-specific substitution in TRPV3 (N647D), a calcium permeable thermoTRP expressed in sensory neurons (Smith et al., 2002), skin (Xu et al., 2002) and epidermal keratinocytes (Mandadi et al., 2009), and hair follicles (Cheng et al., 2010) that functions as a warm temperature sensor (>33°C) and regulator of hair growth (Cheng et al., 2010; Smith et al., 2002; Xu et al., 2002), occurred in the pore loop (**Fig. 7A/B**) and was fixed for the derived aspartic acid in seven woolly mammoth individuals we tested by PCR amplification and Sanger sequencing (**SI Fig. 6**). Remarkably, a previous high-throughput mutagenesis screen found that mutations at site 647 abolished heat sensitivity in mouse TRPV3 (Grandl et al., 2008), suggesting the N647D substitution affects thermosensation by mammoth TRPV3.

**Figure 7.**
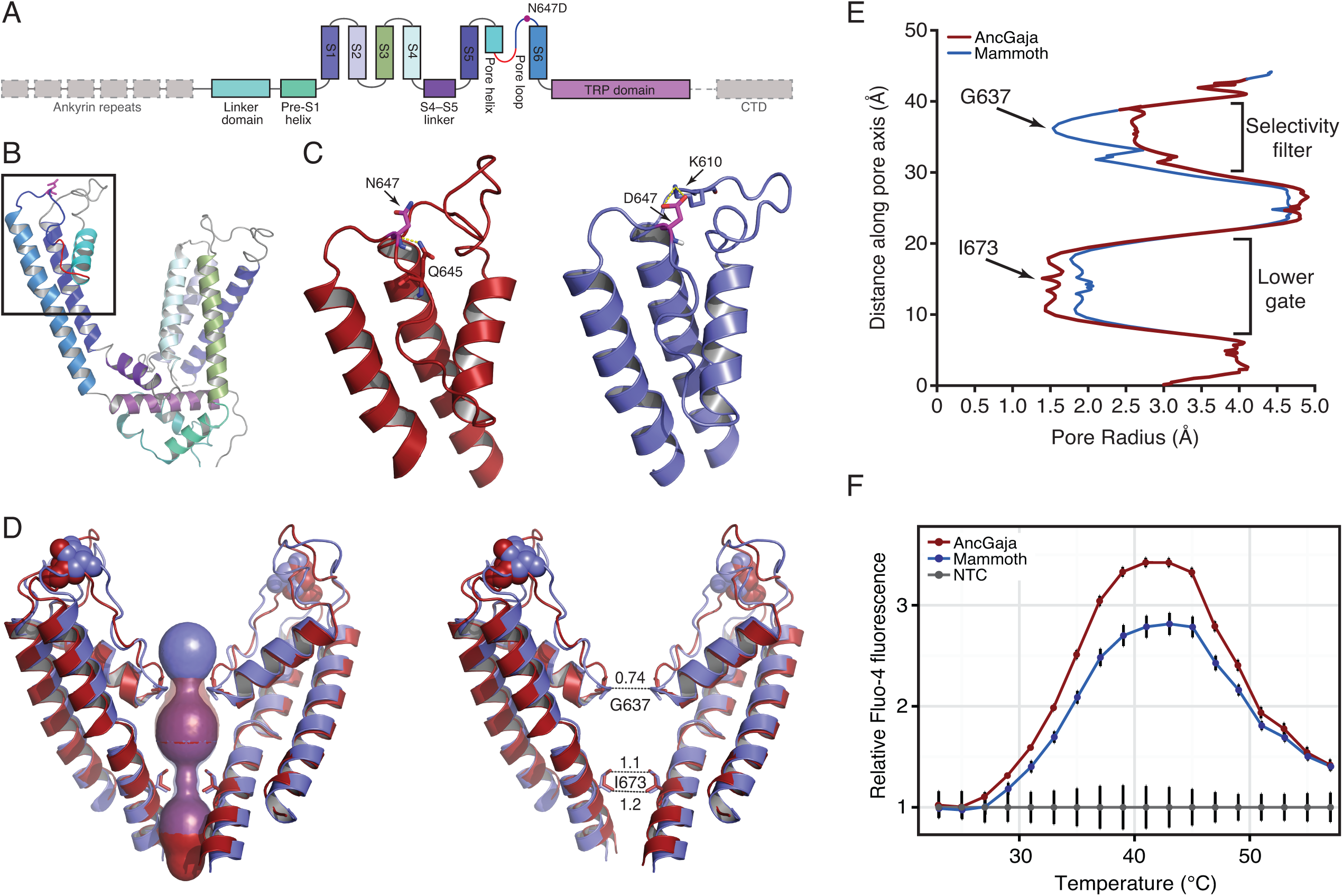
Structural and functional consequences of the Woolly mammoth-specific N647D substitution in TRPV3. (A) Diagram of the major structural domains of TRPV3. Grey regions were not included in the TRPV3 structural model. The location of the mammoth-specific N647D substitution is shown as a magenta circle and the selectivity filter as a red loop. (B) Cartoon representation of the pore domain of the TRPV3 homology model. The N647D substitution is shown in stick representation and colored magenta and the selectivity filter as a red loop. The region shown in panel (C) is boxed. (C) Close up view of the pore region of the AncGajah (red) and AncMammoth (blue) TRPV3 homology models. N647 in AncGajah and D647 in AncMammoth are colored magenta and shown in stick representation, predicted hydrogen bond interactions with neighboring residues are shown as yellow dashed lines. (D) Superimposed AncGajah (red) and AncMammoth (blue) pore regions in the open conformation. Only diagonally opposes subunits of the shown. Site 647 is shown as spheres, sites G637 and I673 are sticks. Left, predicted pores formed by the AncGajah (red) and AncMammoth (blue) TRPV3 channels are shown as space filling spheres. Right, diameter of the AncMammoth pore relative to the diameter AncGajah pore at the narrowest point in the selectivity filter (site G637) and lower gate (site I673). (E) Profile of the predicted pore radius from AncGajah (red) and AncMammoth (blue) TRPV3 channels. (F) Flour-4 fluorescence intensity in response to increases in temperature in HEK293 cells transiently transfected with expression constructs for AncGajah (red) and AncMammoth (blue) TRPV3 relative to non-transfected cells (grey). Curves shown as background subtracted relative intensity, mean±SEM (n=6).

We inferred the structural consequences of the N647D substitution by generating homology models of the AncGajah and AncMammoth TRPV3 protein and tetrameric channel (as desceibed above). We found that the TRPV3 models were structurally very similar to the TRPV1 reference structure on which they were based (RMSD = 1.93-1.99 Å **SI Fig. 7**), particularly in the α-helices that form the pore of the tetrameric channel suggesting these are realistic models. In the TRPV1 structure, hydrogen bonds between residues in the pore loop are thought to maintain the outer pore in a non-conductive conformation in the closed state; conformational changes in the pore helix and pore loops disrupt these local hydrogen bonds to facilitate gating and widening of the selectivity filter upon channel activation (Cao et al., 2013; Liao et al., 2013). Our structural model suggests that the carbonyl oxygen of the ancestral N647 residue forms a hydrogen bond with the neighboring side chain of Q645, whereas in the AncMammoth structure these hydrogen bonds are replaced by a pair of hydrogen bonds between D647 and K610, likely impeding full opening of the channel in mammoths (**Fig. 7C**). Indeed in the model of the AncGajah open channel the distance between diagonally apposed G637 residues is 8.5 Å whereas this distance is only 6.3 Å in the AncMammoth open channel (**Fig. 7D/E**), which narrows the pore diameter at the selectivity filter by ∼26% and the pore radius at the selectivity filer by ∼60% (**Fig. 7D/E; SI Fig. 8**).

To functionally characterize the effects of the mammoth-specific N647D substitution we resurrected the AncMammoth and AncGajah TRPV3 genes and measured their temperature-dependent gating in transiently transfected human embryonic kidney cells (HEK293) using Fluo-4 calcium flux assays (Aneiros and Dabrowski, 2009; Reubish et al., 2009). We found that both the AncMammoth and AncGajah TRPV3 proteins were expressed at similar levels (**SI Fig. 8**) and that the overall gating dynamics of channels were very similar. Both channels, for example, are activated at ∼29°C, had half maximum activities (T_50_) at 33°C, and maximal activities (T_max_) at 43°C (**Fig. 7F**). The AncMammoth TRPV3 channel, however, was ∼20% less active at than the AncGajah channel at T_max_ (**Fig. 6F**), consistent with the predictions from our structural models that the AncGajah channel does not fully open upon stimulation. We found that neither the AncMammoth nor AncGajah channels were activated by 2-APB, however, while both channels were robustly activated in response to camphor (**SI Fig. 8**). These data are consistent with previous studies in mice that found mutations at site 647 affect temperature-dependent gating but not channel opening by chemical agonists (Grandl et al., 2008).

To test whether this substitution may have been positively selected in the woolly mammoth lineage, we assembled a dataset of *TRPV3* genes from 64 diverse amniotes and used maximum likelihood methods to identify lineages (aBSREL) and codons (MEME) with evidence of episodic and pervasive (FEL) diversifying selection. While aBSREL identified a class of sites in mammoth with *d*_*N*_*/d*_*S*_>1, the results were not significant (mean *d*_*N*_*/d*_*S*_=10, *P*=0.226). MEME, however, found significant evidence for episodic diversifying selection at site 647 (*d*_*N*_*/d*_*S*_=444.47, *P*=0.037); although inferences of positive selection at specific branch-site combinations are inherently imprecise, the MEME model suggested the N647D substitution was positively selected in mammoths (PP>0.97, EBF>1000). In contrast FEL inferred site 647 to evolve under strong purifying selection (*d*_*N*_*/d*_*S*_=0.274, *P*=0.044) indicating this site does not experience pervasive diversifying selection. These data suggest that while *TRPV3* genes and site 647 generally evolve under purifying selection, there is strong evidence that the N647D substitution was positively selected in stem-lineage of woolly mammoths.

Our observation that the mammoth TRPV3 protein is less active (hypomorphic) across a range of temperatures is particularly intriguing given its dual roles in temperature sensation and hair growth. *TRPV3* knockout mice, for example, have deficits in responses to innocuous and noxious heat and prefer colder temperatures than wild-type mice (Marics et al., 2014; Miyamoto et al., 2011; Moqrich et al., 2005). TRPV3 activation also inhibits hair shaft elongation and induces the premature regression hair follicles (Borbíró et al., 2011; Cheng et al., 2010), whereas *TRPV3* knockout mice have curly whiskers and wavy hair (Cheng et al., 2010). These data suggest that the hypomorphic mammoth TRPV3, which partly phenocopies *TRPV3* null mice, may have directly contributed to evolution of cold tolerance and long hair in mammoths.

## CONCLUSIONS

Identifying the genetic changes that underlie morphological evolution is challenging, particularly in non-model and extinct organisms. We have identified genetic changes unique to woolly mammoths, some of which likely contributed to woolly mammoth-specific traits. Our results suggest that changes in circadian systems, insulin signaling and adipose development, skin development, and temperature sensation may have played important roles in the adaptation of woolly mammoths to life in the high artic. Our identification of a hypomorphic woolly mammoth amino acid substitution in TRPV3 is particularly noteworthy given its dual roles in temperature sensation and hair growth, suggesting that this substitution played a direct role in the origin of extreme cold tolerance.

## EXPERIMENTAL PROCEDURES

### Genome sequencing, assembly, and annotation

Details of the sequencing protocol are given under Extended Experimental Procedures. Sequences from the three Indian elephant samples were aligned to the reference genome from the African Elephant (loxAfr3) using the Burrows Wheeler Aligner (Li and Durbin, 2010) with default parameters (BWA version 0.5.9-r16). The reads were subsequently realigned around putative indels using the GATK (DePristo et al., 2011) IndelRealigner (version 1.5-21-g979a84a), and putative PCR duplicates were flagged using the MarkDuplicates tool from the Picard suite (version 1.96).

For the two mammoth samples, we trimmed putative adapter sequences and merged overlapping paired-end reads using available scripts (Kircher, 2012). We required an overlap of at least 11 nucleotides between the mates, and only pairs that could be merged were retained for subsequent analyses. The merged reads were aligned to the genome from the African elephant (loxAfr3) using BWA with default parameters, and only the mapped reads that were longer than 20 bps were retained for the subsequent SNP calls. The reads were realigned using the GATK IndelRealigner and putative PCR duplicates were flagged using MarkDuplicates, similar to the process described for the modern genomes. We also limited the incorporation of damaged sites into the variant-calling pipeline by hard-masking all sites that would be potentially affected by the characteristic ancient DNA patterns of cytosine deamination in single stranded overhangs. This mask was applied to 10 nucleotides on both ends of the merged reads from the ancient samples.

At about 33 million positions in the African elephant reference assembly we detected a nucleotide different from the reference in at least one of the five newly sequenced individuals. We call these positions single-nucleotide variants (SNVs); these were identified using SAMtools (Li et al., 2009) (version 0.1.19), which was applied with “-C50” to adjust the mapping quality of the reads with multiple mismatches. We did not call differences in regions where the reference base was unknown, and the calls were limited to regions that were covered at least 4 times, and at most 250 times by the sequences in these samples.

We selected the SNVs where the two mammoths were identified as homozygous for the variant nucleotide, whereas the three Asian elephants were homozygous for the *Loxodonta africana* reference nucleotide. Since the African elephant is thought to have diverged from the ancestor of Asian elephants and mammoths (Krause, et al., 2006), we considered the “fixed” mammoth variant as derived (i.e., non-ancestral). We used the gene annotation for *Loxodonta africana* to identify putative variant amino acids. This information as well as GO terms and gene models were obtained from the ENSEMBL database (Flicek et al., 2013).

We also wanted to provide each SNV with “quality values” that can help determine the robustness of an analysis to potential erroneous SNV calls. It was not clear to us how to define a single quality value that treats mammoths on an equal footing with Asian elephants, because of the lower coverage, shorter length and decreased accuracy of the mammoth reads, so we annotate each SNV with a “mammoth quality value” and an “Asian elephant quality value”, which give the Phred scaled probability of the alternate allele in the two mammoth samples and the three Asian elephants, respectively. Slightly under half of the ∼33 million SNV calls have a mammoth quality value of at least 100, but of the 2,046 putative fixed mammoth-specific non-synonymous differences (2,020 amino-acid variants and 26 premature stop codons), 1,975 have mammoth quality value at least 100, which suggests to us that our conclusions are reasonable robust. In any case, the user can filter the putative SNVs as desired.

A table of all 2,046 fixed, mammoth-specific protein differences is freely available on the Galaxy server (Goecks et al., 2010; Bedoya-Reina et al., 2013; usegalaxy.org). The table has the following columns: 1) gene name; 2) reference amino acid; 3) position in the peptide sequence (base 1); 4) variant amino acid; 5) name of Ensembl transcript; 6) name of scaffold in the *Loxondonta* genome assembly; 7) position in the scaffold (base 0); 8) name of orthologous human chromosome; 9) human position; 10) BLOSUM80 exchangeability score; 11) PolyPhen-2 category (“benign”, “possibly damaging”, “probably damaging”, “unknown”); 12) PolyPhen-2 score; 13) mammoth SNV quality value. Tables of the 33 million SNVs, *Loxodonta*/Ensembl-annotated genes, 170,274 SNVs in those protein-coding regions, and a complete command history for constructing the table of 2,046 differences (see above) are available at http://usegalaxy.org/r/woolly-mammoth).

### Functional inference of mammoth-specific amino acid substitutions

We used VLAD (http://proto.informatics.jax.org/prototypes/vlad/) to mine the mouse knockout (KO) phenotype data at Mouse Genome Informatics (http://www.informatics.jax.org) for the genes with mammoth-specific substitutions. Enriched Gene Ontologies (GO) and KEGG pathways were identified with WebGestalt (http://bioinfo.vanderbilt.edu/webgestalt/). The results can be found in the supplementary file “Enrichment tables.xlsx”.

### TRPA1 and TRPV4 structure modeling

The AncMammoth and AncGajah TRPA1 and TRPV4 protein structures were modeled using the recently published high-resolution cryo-EM structure of TRPV1 in the closed and open states (Cao et al., 2013; Liao et al., 2013). Initial structural models of the AncMammoth and AncGajah proteins in the open and closed were generated using I-TASSER (Roy et al., 2010; Zhang, 2008) and the experimentally determined structure of the TRPV1 channel in the closed (Protein Database accession number 3J5P) and open state (Protein Database accession number 3J5Q) as a templates. Initial AncMammoth and AncGajah structure models were refined with ModRefiner (Xu and Zhang, 2011) using the TRPV1 channel in the closed (PBD ID: 3J5P) and open state (PBD ID: 3J5Q) as the reference structure.

### TRPV3 ancestral sequence reconstruction and gene synthesis

To reconstruct the mammoth ancestral TRPV3 protein sequences we: 1) included TRPV3 genes from the genomes of two woolly mammoths, three Asian elephants, African elephant (loxAfr3), West-Indian manatee, hyrax (proCap1), lesser hedgehog tenrec (TENREC), and nine-banded armadillo (dasNov2); 2) aligned the translated sequences with MUSCLE (Edgar, 2004a,b); 3) inferred the JTT model as best-fitting (AIC=7255.71, cAIC=7256.05, BIC=7380.52) model of amino acid substitution using the model selection module implemented in Datamonkey (Delport et al., 2010); and 4) used joint (Pupko et al., 2000), marginal (Yang et al., 1995), sampled (Nielsen, 2002) maximum likelihood methods implemented in the ASR module of Datamonkey incorporating a general discrete model of site-to-site rate variation with 3 rate classes, using the following phylogeny:

((((((mammoth, Asian elephant) African elephant) manatee) hyrax) tenrec) armadillo)

We found that support for the reconstructions at site 647 in both ancestral sequences was 1.0 under joint, marginal, and sampled likelihoods.

The Asian elephant/mammoth (AncGajah) ancestral TRPV3 gene, including an amino terminal FLAG tag (MDYKDDDDK) with a Kozack sequence incorporated into the FLAG tag (gccaccATGG) and a 4 residue glycine spacer (GGGG) amino (N)-terminal to the TRPV3 open reading frame, was synthesized by GeneScript using human codon-usage, and cloned into the mammalian expression vector pcDNA3.1(+) (Invitrogen). The ancestral mammoth gene (AncMammoth) was generated by using site directed mutagenesis to introduce the N647D mutation into the AncGajah TRPV3 pcDNA3.1(+) construct. The sequence of both ancestors was verified with Sanger sequencing.

### TRPV3 function assays

We used the Fluo-4 NW Calcium Assay Kit (Life Technologies) to determine the temperature-response of the AncMammoth and AncGajah TRPV3 proteins. Human embryonic kidney cells, HEK293 cells (ATCC CRL-1573), were cultured in MEM supplemented with 10% (v/v) FBS in a 37°C humidity controlled incubator with 10% CO_2_. HEK293 cells growing in 10-cm plates were transiently transfected at 80% confuency with 24 μg of expression vector for either the AncMammoth or AncGajah TRPV3 genes, or empty pcDNA3.1(+) using Lipofectamine LTX+ (Life Technologies) using the standard procotol.

48 hours after transfection, cells were harvested by trypsinization, centrifuged, resuspended in Hanks Balanced Salt Solution (HBSS) and HEPES assay buffer containing 2.5mM probenecid, and transferred to a 96-well plate (at 150,000 cells/well in 50ul of assay buffer). Temperature dependent calcium influx was assayed using the Fluo-4 NW Calcium Assay Kit (Molecular Probes) and a high-throughout qPCR-based assay (Aneiros and Debrowski, 2009; Reubish et al., 2009). After an initial 30 min loading at 25°C the temperature was raised from 15°C to 57°C in 2°C steps and fluorescence measured after 2 minutes at each temperature using a BioRad CFX-96 real time PCR machine. Fluo-4 fluorescence was measured using channel 1 (Sybr/FAM). Fluo-4 fluorescence of cells transfected with the AncMammoth or AncGajah TRPV3 genes was normalized by the Fluo-4 fluorescence of empty pcDNA3.1(+) transfected controls. All experiments included 6 biological replicates, and were repeated in 4 independent experiments.

### TRPV3 structure modeling

The AncMammoth and AncGajah protein structures were modeled using the recently published high-resolution cryo-EM structure of TRPV1 in the closed and open states (Cao et al., 2013; Liao et al., 2013). Initial structural models of the AncMammoth and AncGajah proteins in the open and closed were generated using I-TASSER (Roy et al., 2010; Zhang, 2008) and the experimentally determined structure of the TRPV1 channel in the closed (Protein Database accession number 3J5P) and open state (Protein Database accession number 3J5Q) as a template. Initial AncMammoth and AncGajah structure models were refined with ModRefiner (Xu and Zhang, 2011) using the TRPV1 channel in the closed (PBD ID: 3J5P) and open state (PBD ID: 3J5Q) as the reference structure. The backbone atoms of the refined AncMammoth and AncGajah structure models in their closed and open conformations were then aligned to the pore tetramer of the TRPV1 structure.

## AUTHOR CONTRIBUTIONS

VJL designed and led the experimental analysis of TRPV3, VJL, WM and OCBR analyzed the elephantid sequences, AR identified sequence variants, MS performed experiments on TRPV3, DIM performed PCR validations, GP provided insights about sequence analysis and experimental methods, SCS and WM led the woolly-mammoth sequencing project, VJL and WM wrote the paper with input from the coauthors.

## DATA AVAILABILITY

The Illumina reads from the five sequenced elephantid genomes are available in the Short Read Archive, accession numbers have not been assigned. Tables of the nucleotide and amino-acid differences that we identified and a table of putative gene gains and losses are available at the Galaxy website, and collected at http:usegalaxy.org/r/woolly-mammoth, along with the table of putative fixed woolly mammoth-specific amino acids and the set of Galaxy commands that created it. Those data can be further analyzed by a suite of Galaxy tools designed specifically for these datatypes (Bedoya-Reina et al., 2013)

## ACKNOWLEDGEMENTS

Sequencing was supported by NSF award EAR-0921958 to SCS. We thank O. Ryder and S. Santosh for providing Asian elephant DNA samples, and M. DeGiorgio for insights about woolly-mammoth SNVs. We also thank S. L. Kosakovsky Pond for assistance in interpreting the TRPV3 selection tests results.

## Extended Experimental Methods

### Sequencing

We greatly expanded the sequence coverage of two samples that we had analyzed earlier (Miller et al., 2008). DNA from our M4 and M25 hair samples was extracted following the previously described protocol (Gansuage and Meyer, 2013).

In total, 13 different libraries were constructed for mammoth M25 (internal IDs 7, 7B, 7new, 7bead, 109B, 109C, 110B, 110C, 130A, 131A, 131C, 164 and 165). Libraries 7 and 7new were prepared using Illumina’s Genomic DNA Library Preparation Kit, starting with 800ng of DNA. The remaining libraries were prepared with a hybrid approach for which Roche 454 Rapid Library Preparation reagents were used in combination with Illumina DNA adapters. The finished libraries were then subjected to Roche 454 emulsion PCR instead of the standard Illumina pooled PCR. The intent of this approach was to overcome the PCR inhibiting effects of melanin, which is always coextracted together with DNA when isolated from hair.

Library 7 was sequenced in five lanes of Illumina GA II paired-end sequencing runs with varying read lengths of 36-42 basepairs (bp), generating a total of 99,281,792 reads or 5,249,221,008 bases. In addition, library 7 was also subjected to 22 lanes of Illumina GA IIx paired-end sequencing at varying read lengths of 76-82 bp, generating an additional 1,394,299,608 reads or 110,611,021,888 bases.

Library 7B was sequenced in one lane on a 80×80 bp paired-end Illumina GA IIx sequencing run, generating 65,689,518 reads or 5,255,161,440 bases. Library 7new was sequenced on both, the Illumina GA IIx as well as the Illumina HiSeq2000 platforms, at read lengths of 82 and 101 bp, respectively. In total, 979,063,102 reads or 80,283,174,364 bases were generated on the GA IIx for this library and 2,079,562,938 reads or 210,035,856,738 bases on the HiSeq2000 platform.

The Roche/Illumina hybrid libraries were each sequenced in one lane on the Illumina GA IIx sequencing platform at a read length of 82 bp paired-end (library 7bead), 76 bp paired-end (libraries 110B, 164, and 165) or 36 bp single-read (libraries 109B, 109C, 110B, 110C, 130A, 131A, and 131C), generating a combined number of reads of 452,525,020 or 28,471,674,736 bases.

For mammoth M4, 9 different libraries were constructed (internal IDs 17, 288B, 289B, 290B, SCELSE215, SCELSE491A-C, SCELSE492A-C, SCELSE493A-C, SCELSE494A-C). Library 17 was prepared using Illumina’s Genomic DNA Library Preparation Kit, starting with 100ng of DNA. Libraries 288B, 289B, and 290B were prepared with a hybrid approach, using Roche 454 Rapid Library Preparation reagents in combination with Illumina DNA adapters. PCR-based enrichment of the finished libraries was then performed according to Illumina’s amplification protocol. Library SCELSE215 was prepared with the Illumina TruSeq DNA Library Preparation Kit, following the manufacturers’ recommendations. Libraries SCELSE491A-C, SCELSE492A-C, SCELSE493A-C, SCELSE494A-C were prepared following the single-stranded DNA library preparation protocol^25^ that was specifically developed for ancient or damaged DNA, which typically does not perform well with commercially available library preparation kits. In short, DNA samples (10pg-10ng) are dephosphorylated and heat denatured to create single-stranded fragments to which a single-stranded, biotinylated adapter is ligated. The ligation product is then immobilized on streptavidin beads and the second strand is filled in by primer extension. 3’ overhangs are removed to create blunt-end fragments, a second adapter is ligated to the library which is then eluted off the beads. A PCR optimization step is performed to determine the optimal PCR cycle number. The library is then amplified using the previously determined optimal PCR cycle number and a unique combination of index primers for each sample, which allows for library pooling during sequencing.

Performing library preparation for ancient DNA samples at the single-stranded DNA level has the following advantages:

1. Ancient DNA molecules are typically of very short length (<100bp) and are often lost in purification steps that utilize silica spin columns or carboxylated beads. Biotinylating the ancient DNA molecules will allow for all library preparation and purification steps to be carried out while the DNA is tightly bound to streptavidin beads, therefore avoiding the loss of molecules.
2. Ancient DNA is often heavily degraded and many DNA fragments contain single-strand breaks. These molecules will be lost with double-stranded DNA library preparation methods whereas the single-stranded DNA library preparation method disassembles these molecules into multiple independent fragments upon heat denaturation with each fragment potentially contributing to the sequencing library.
3. Nucleotide modifications located at the end of one of the two complementary DNA strands may inhibit adapter ligation during double-stranded DNA library preparation. With the single-stranded method, the unmodified strand can still be retrieved.

The M4 single-stranded DNA libraries were prepared with 5ng, 15ng, 5ng, and 5ng of DNA, respectively. The DNA starting material for these libraries originated from 3 distinct DNA extractions. Our protocol^4^ was used for the extractions as described above with the following modification: For the DNA extraction that was used for libraries SCELSE491A-C and SCELSE492A-C, the mammoth hair was decontaminated with 5% bleach for 1 minute, followed by a water wash prior to DNA extraction. For the DNA extractions that were used for libraries SCELSE493A-C and SCELSE494A-C, the decontamination time with bleach was increased to 5 minutes and 10 minutes, respectively.

Library 17 was sequenced in one lane of an Illumina GA II paired-end sequencing run at a read length of 36 bp, generating 22,081,140 reads or 794,921,040 bases. Library SCELSE215 generated a total of 7,562,918 reads or 1,142,000,618 bases from one Illumina MiSeq 151x151 bp paired-end sequencing run. Libraries 288B, 289B, and 290B were sequenced in a total of 17 lanes of Illumina HiSeq2000 paired-end runs at a read length of 75-101 bp, generating 4,705,377,806 reads or 458,623,900,478 bases. In addition, libraries 288B, 289B, and 290B were also sequenced in one Illumina MiSeq single-read run at a read length of 50bp. This run generated 2,146,802 reads or 107,340,100 bases. A total of 2,340,774,696 reads (175,558,102,200 bases) were generated from libraries SCELSE491A-C, SCELSE492A-C, SCELSE493A-C, and SCELSE494A-C which were sequenced in 8 lanes of HiSeq2500 rapid sequencing runs at a read length of 75 bp paired-end.

In addition to the 1,076 Gb (billion bases) that were generated for the mammoth samples M4 and M25 on the Illumina platforms, three Indian elephants were also sequenced to generate an “outgroup” species to allow sequence differences between the woolly mammoth and African elephant to be assigned to a particular lineage. For each of the elephant samples (Uno, Parvathy, and Asha), one library was prepared with either the Illumina Genomic DNA Library Preparation Kit (Uno, internal ID 186B) or the Illumina TruSeq DNA Library Preparation Kit (Parvathy, internal ID SCELSE8A and Asha, internal ID SCELSE9A). Library 186B was sequenced in 4 lanes of Illumina GA IIx runs at a read-length of 82 bp paired-end and in 6 lanes of Illumina HiSeq2000 runs at a read-length of 101 bp paired-end. Libraries SCELSE8A and SCELSE8B were each sequenced in 4 lanes of an Illumina HiSeq2000 101 bp paired-end sequencing run. In total, 1,349,450,838 reads or 130.6 Gb (billion bases) were generated for the Indian elephant samples.

### PCR validation of the TRPV3 mammoth-specific N647D substitution

The TRPV3 variant was validated by PCR and Sanger sequencing in 7 different mammoth specimens as well as one Asian Elephant (Asha). The following PCR primer sequences were ordered from IDT Singapore (HPLC purified) and used for PCR validation:

M_P1F: 5’-AGGAGGACGAAGGTGAGGAT-3’

M_P1R2: 5’-CGGTGCTGGAACTCTTCAA-3’

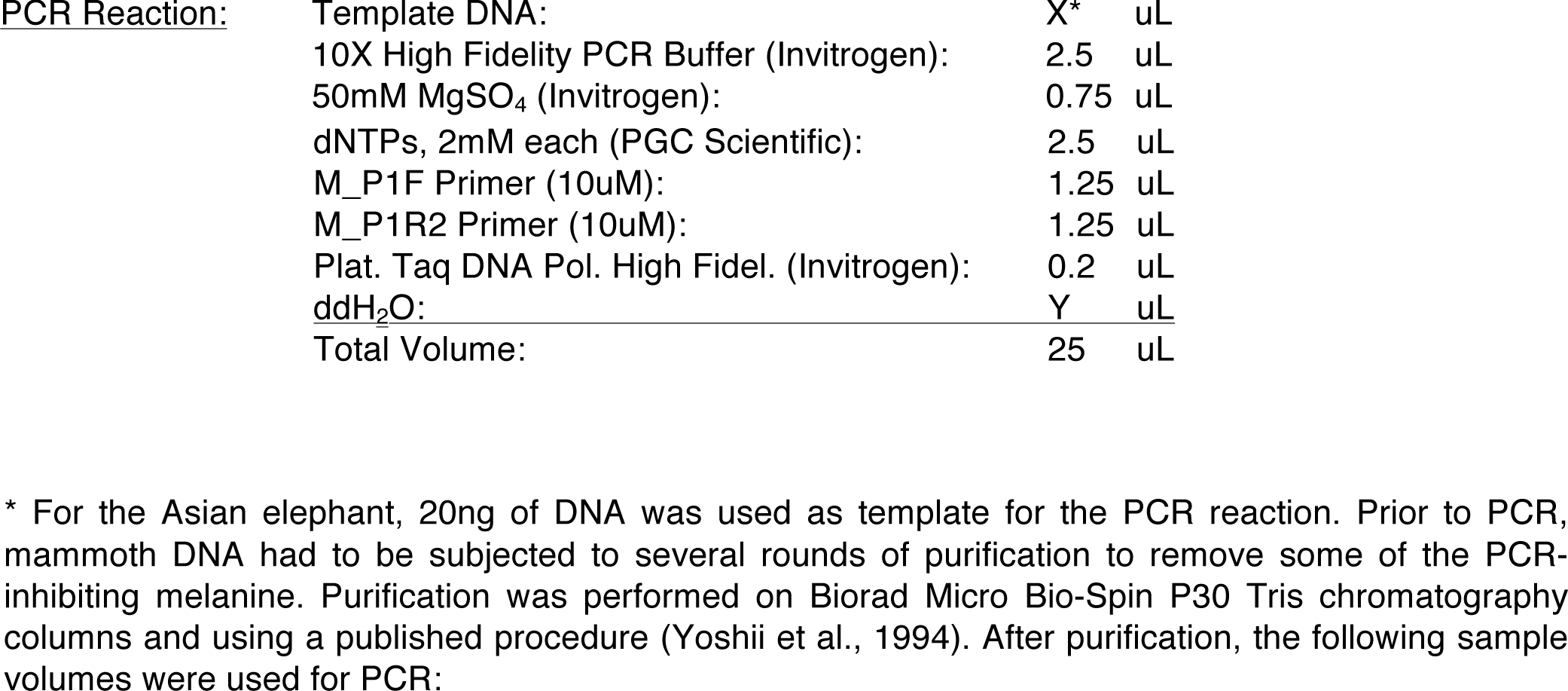

M2: 4 uL

M4: 4 uL

M15: 4 uL

M19: 4 uL

M25: 4 uL

M26: 4 uL

Yukagir: 14 uL

<u>Cycling Conditions (performed on Applied Biosystems GeneAmp 9700):</u>

1. T=94°C; 0:02:00
2. T=94°C; 0:00:20
3. T=53°C; 0:00:20
4. T=68°C; 0:00:30
5. Go to 2. rep 40
6. T=68°C; 0:08:00
7. Hold 4°C

PCR reactions were purified with Agencourt AMPure XP beads (Beckman Coulter) and submitted to AITbiotech, Singapore, for Sanger sequencing. Sanger reads were then aligned to the TRPV3 reference sequence using Sequencher (Gene Codes Corporation). Results are shown in Supplemental Fig. S3.

## Supplemental Information

### Sequencing

The sequencing protocol is described in detail in the Methods section. Fig. S1 summarizes the results.

### Known mammoth protein variants

We looked at genes containing previously studied mutations in mammoth. Fur of different colors has been recovered from permafrost mammoth mummies. A previous study identified three non-synonymous substitutions in the *MC1R* gene of mammoths (i.e. T21A, R67C, and R301S), for which one individual was found to be heterozygous, while the other three were homozygous (Römpler et al. 2006). Orthologs of this gene determine hair color in mammals.

The gene *MC1R* is not annotated in the ENSEMBL gene models for *Loxodonta*, so we identified its position in the genome using the sequence reported by Römpler et al. (2006) in GenBank. We mapped the gene to a unique position in the genome in scaffold 57 between the nucleotides 3,482,559 and 3,483,513. We found none of the published *MC1R* mutations. On the other hand, we found one non-synonymous mutation (E37K) that originated in the mammoth lineage. For this mutation, one individual (M25) seemed to be heterozygous while the other appeared homozygous. This mutation is in the transmembrane domain of the protein, far from the extracellular mutations described to alter the function of the protein.

Another study (Campbell et al., 2010) reported several mutations in the genes encoding the subunits A-T (*HBA-T*) and B/D (*HBB/HBD*) of hemoglobin, and showed that the mutations in the beta/delta subunit decreased the energetic cost of delivering oxygen from lungs. For the *HBA-T* gene the authors did not discovered any polymorphism originated in the mammoth lineage, though they found one non-synonymous mutation originated in the African elephant lineage (S50G), another originated in the mammoth/Asian elephant lineage (A58G), and another in the Asian elephant lineage (N6K). Additionally they found two synonymous mutations in the Asian elephant lineage. We found the mutations A58G and S50G as previously described by Campbell et al. (2010); however the non-synonymous mutation N6K and the synonymous substitutions were absent in our results. In addition, we found one synonymous mutation (L77) heterozygous in the two mammoth individual sequenced, and one non-synonymous mutation (R32W) heterozygous in only one mammoth individual (i.e. M4).

For the beta/delta subunit, the previous study reported three non-synonymous mutations originated in the mammoth lineage (T13A, A87S and E102Q), and another non-synonymous (D53E) and one synonymous in African elephant. We found the amino-acid substitutions reported to be in the African elephant lineage, while only the substitution A87S was found to be originated (and heterozygous) in mammoth individual M4. Nevertheless, we found additional non-synonymous mutations originated in mammoths: Q11L, G17D, and E122A. Q11L and G17D were found to be heterozygous in both mammoth individuals sequenced, while E122A was found to be heterozygous in M4 and homozygous in M25.

### Inter-species conservation at the TRPV3-variant site

The preliminary analyses that justified the effort required to experimentally investigate the TRPV3 variant included checking the evolutionary conservation of the SNV site, as shown in Fig. S2.

### PCR validation of the TRPV3 variant

Another critical test that justifies the experimental analysis of the TRPV3 variant was the use of PCR amplification and sequencing, as described in the Methods section and with results depicted in Fig. S3.

### TPRV3

Many aspects of our close analysis of the TRPV3 variant are displayed in Fig. 7 of the main paper. Additional details are shown in Figs. S4 and S5.

---

**Figure S1.**
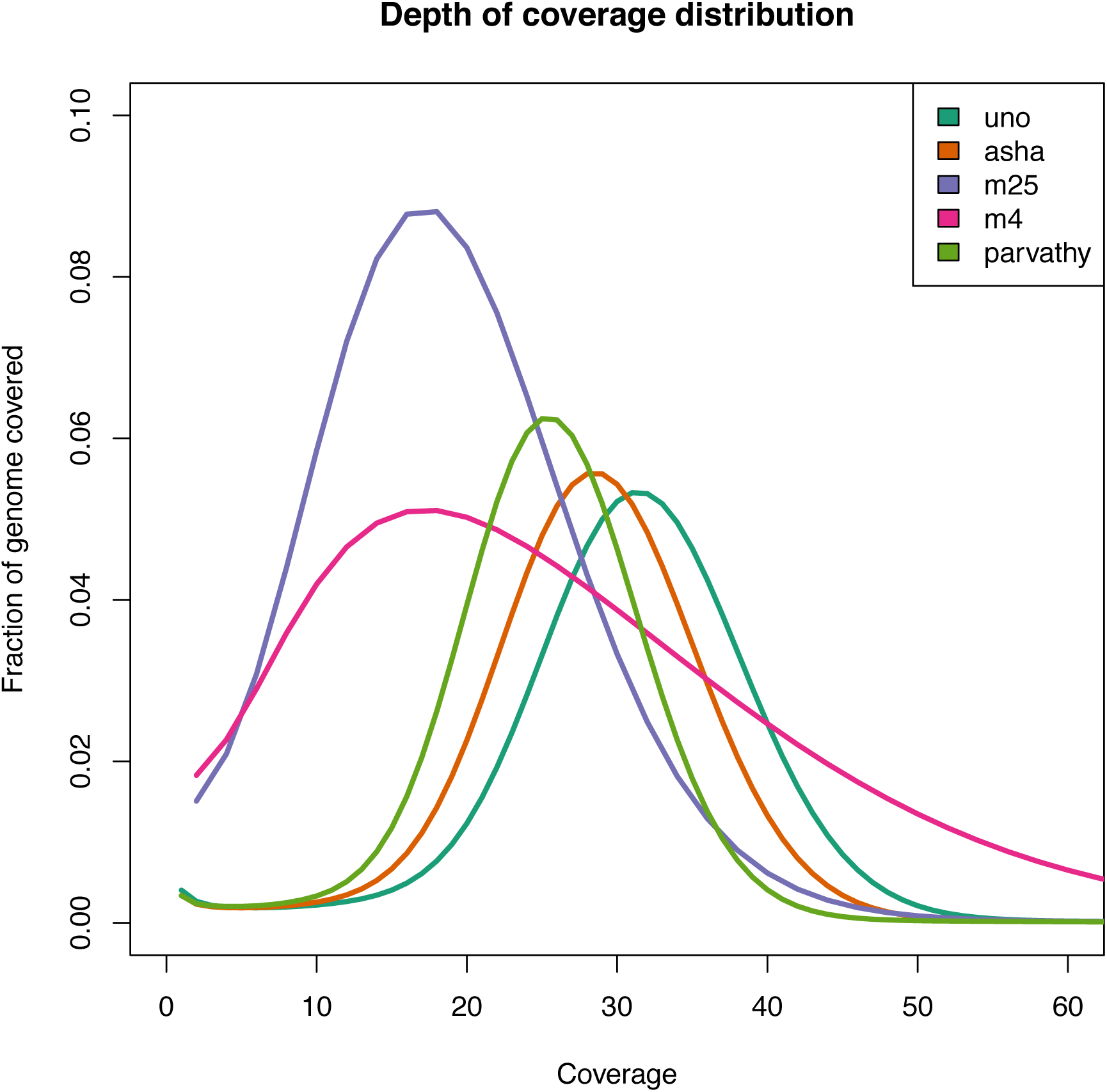
Depth of sequence coverage. The non-redundant reads that aligned uniquely to the African elephant genome had an average coverage of around 20-fold for the two mammoth specimens, M4 and M25, and around 30-fold for the each of the three Asian elephants.

**Figure S2.**
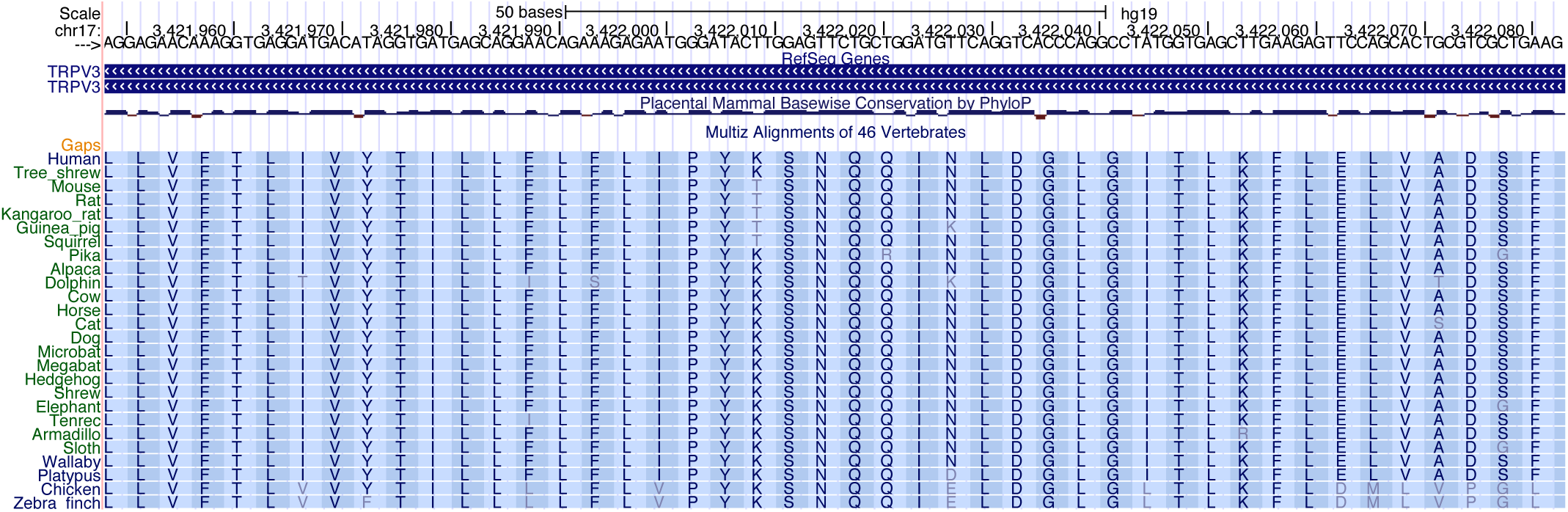
Evolutionary invariance of the TRPV3 region. The figure, created by the UCSC genome browser (http://genome.ucsc.edu; Kent et al., 2002), indicates that the N mutated to D in woolly mammoth is essentially invariant in mammals and birds (center column of the figure). It is in a conserved SNQ motif and, if rodents are omitted, in a conserved PYKSNQQI motif.

**Figure S3:**
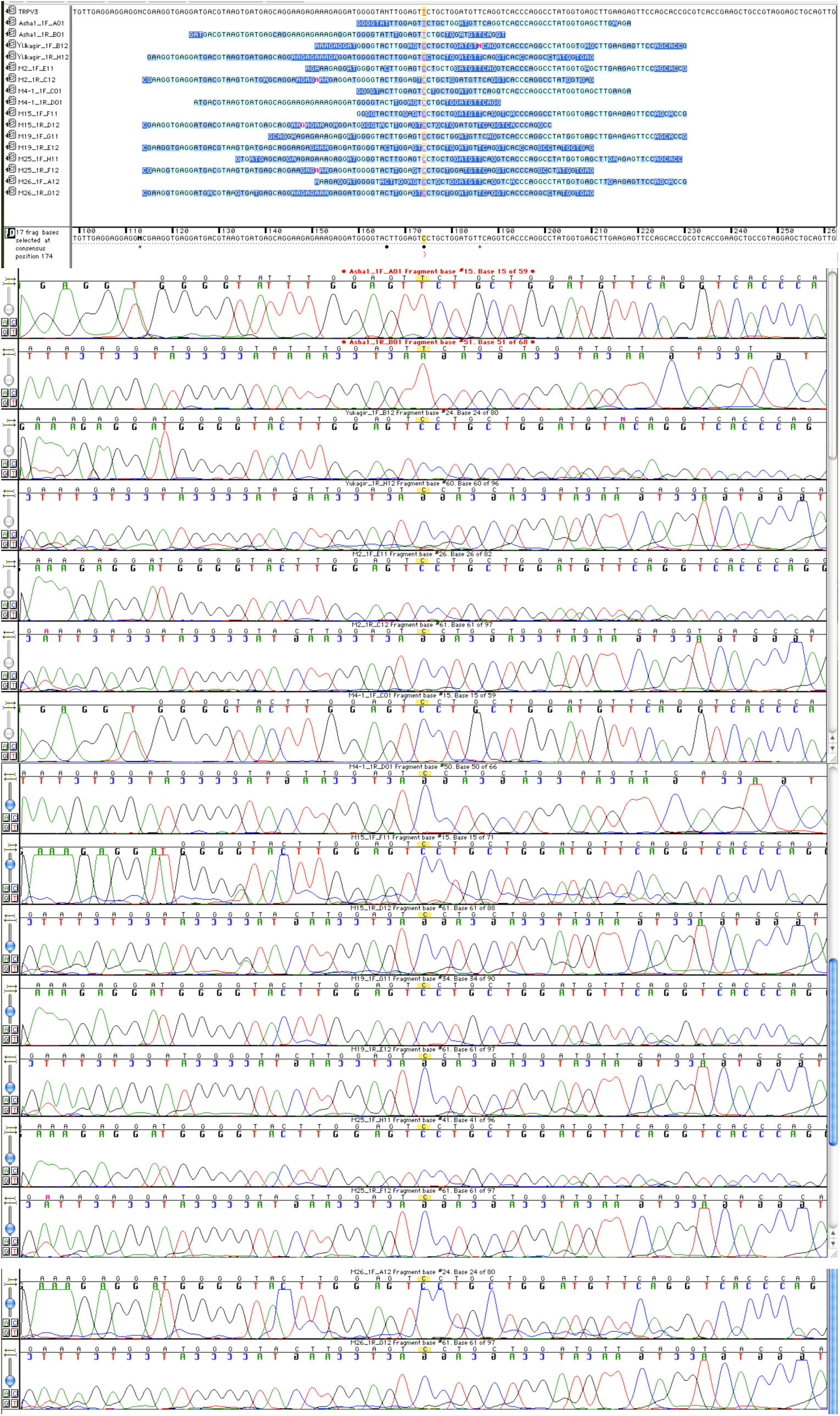
Alignment of the Sanger reads to the TRPV3 reference using Sequencher. The figure shows that both, forward and reverse PCR sequences for the Asian elephant (Asha) are in agreement with the reference sequence at the SNP position (“T”) whereas all mammoth sequences (M2, M4, M15, M19, M25, M26, Yukagir) show a “C” at that position.

**Figure S4.**
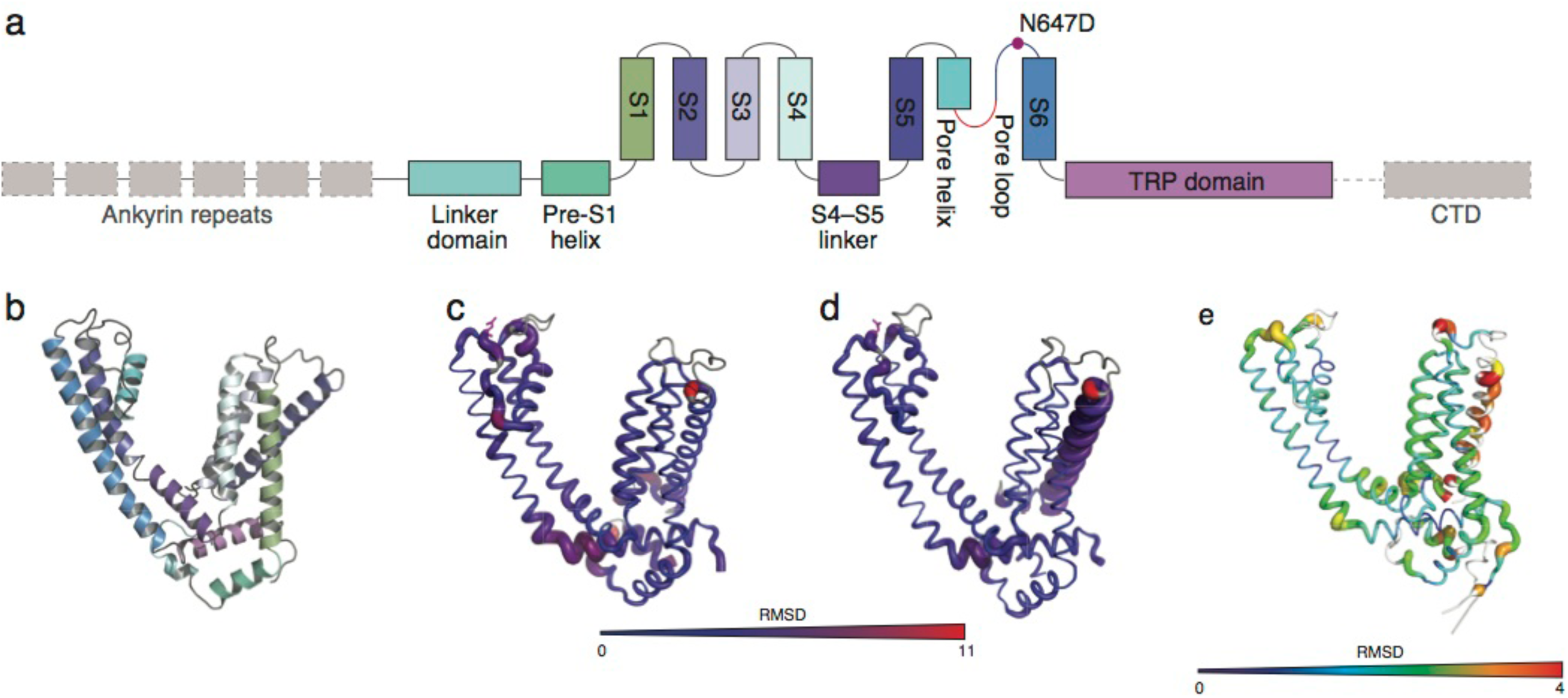
TRPV3 structure models. **a**, Linear diagram showing the major structural domains of TRPV channels. Structural features are colored to match panel b. Dashed grey boxes indicate regions not included in the TRPV3 structural model. **b**, Cartoon representation of the pore domain of TRPV1 reference structure for the the mammoth (AncMammoth) and Asian elephant/mammoth common ancestor (AncGaja) TRPV3 homology models. **c, d,** Putty representation of the pore domain of AncGaja (c) and AncMammoth (d) TRPV3 proteins in the open conformation. The diameter and color of the cartoon are scaled to the RMSD compared to the TRPV1 reference structure in the open state conformation. **e,** Putty representation of the pore domain of AncMammoth TRPV3 homology model in the open conformation. The diameter and color of the cartoon are scaled to the RMSD compared to the AncGaja TRPV3 homology model in the open conformation

**Figure S5.**
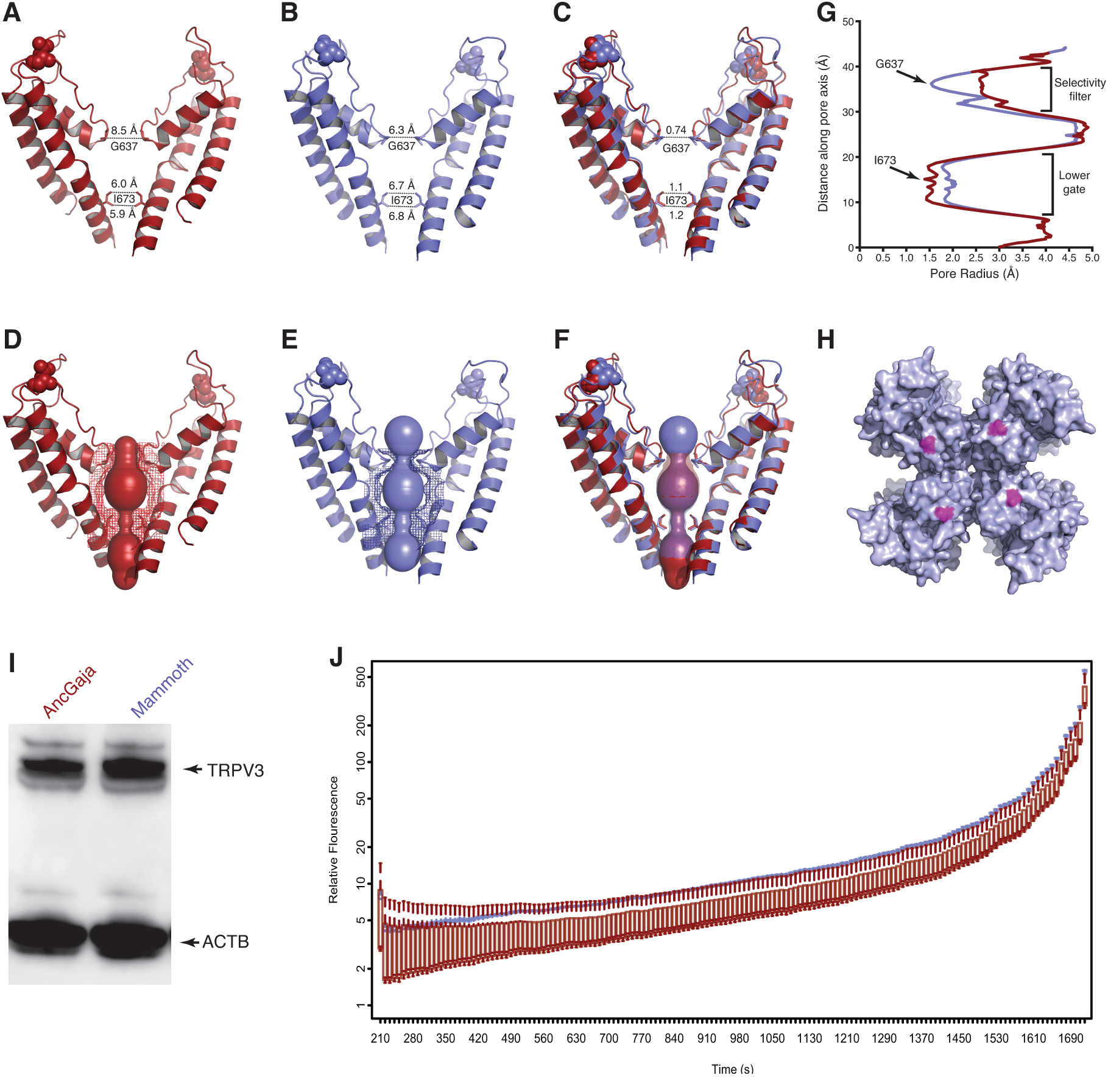
Structural and functional consequences of the mammoth-specific N647D substitution in TRPV3. **a-c**, Cartoon representation of the pore domain of the TRPV3 homology models in the open conformation. Only diagonally opposes subunits of the shown. Site 647 is shown as spheres, sites G637 and I673 are sticks. **a**, AncGaja (red). **b**, AncMammoth (blue). **c**, Superimposed AncGaja and AncMammoth pore TRPV3. **a**, **b**, diameter of the AncGaja and AncMammoth pores at the narrowest point in the selectivity filter (site G637) and lower gate (site I673). **c**, diameter of the AncMammoth pore at sites G637 and I673 relative to the AncGaja pore. **d-f**, predicted pores formed by the AncGaja (d) and AncMammoth (e) TRPV3 channels are shown as space filling spheres. **f**, Superimposed AncGaja (red) and AncMammoth (blue) pore regions. **g**, profile of the predicted pore radius from AncGaja (red) and AncMammoth (blue) TRPV3 channels. **h**, location of site 647 on the extracellular surface of TRPV3 (pore is at the center). **i,** expression of resurrected AncGaja and AncMammoth TRPV3 proteins in HEK293 whole cell lysates. **j**, Flour-4 fluorescence intensity over time (seconds) in response to camphor in HEK293 cells transiently transfected with expression constructs for AncGaja (red) and AncMammoth (blue) TRPV3 relative to non-transfected cells. Boxplots shown as background subtracted relative fluorescence intensity (n=3).

